# Selective Transport of Plasma-Derived Reactive Species through the Plant Aquaporin Channels: A Molecular Dynamics Study

**DOI:** 10.64898/2025.12.22.695874

**Authors:** Davronjon Abduvokhidov, Parthiban Marimuthu, Akbar Kodirov, Mukhammadali Niyozaliev, Dingxin Liu, Jamoliddin Razzokov

## Abstract

The selective permeability of reactive oxygen and nitrogen species (RONS), generated by cold atmospheric plasma (CAP), through plant aquaporins was investigated to identify plasma-derived species capable of intracellular delivery. Using atomistic molecular dynamics and enhanced sampling methods, we quantified the free energy profiles of eight RONS (HNO_3_, HO_2_, *cis*-HNO_2_, *trans*-HNO_2_, N_2_O_4_, NO, NO_2_, and O_3_) across the PIP2;1 aquaporin channel embedded in a lipid bilayer. Hydrophobic species such as NO and O_3_ exhibited minimal energy barriers (~1-2 kJ·mol^−1^) facilitating rapid permeation, while polar and bulky molecules like HNO_3_ and N_2_O_4_ encounter substantial energy barriers (>15 kJ·mol^−1^), particularly near the selective region (also known as the ar/R constriction), which acts like a filter to control what can pass through the aquaporin. These results reveal that RONS permeability is governed by molecular size, polarity, and hydrogen bonding capacity. This mechanistic insight enables rational selection of CAP-generated species for enhancing plant uptake efficiency, with implications for sustainable plasma-based agricultural technologies.

## Introduction

Recent advances in sustainable agriculture have pointed to cold atmospheric plasma (CAP) technology as a promising eco-friendly technology to enhance seed germination, stimulate plant growth, and improve crop yield [1–3]. CAP produces a mixture of short-lived and long-lived reactive oxygen and nitrogen species (RONS), including H_2_O_2_, HO_2_,·OH, NO, NO_2_, O_3_, OH, N_2_O_4_ HNO_3_, s-*cis*-HONO, and s-*trans*-HONO which are believed to influence plant physiology when applied externally to seeds or tissues [1, 4]. Experimental studies have demonstrated that plasma treatment can accelerate germination rate, early plant development, shorten the ripening period, and reduce irrigation demands - yielding significant agronomic and environmental benefits [5–8]. However, the fundamental molecular mechanisms underlying these effects remain unclear. One hypothesized pathway involves the interaction of plasma-generated RONS with cellular transport systems. In this context, aquaporins (AQPs) have emerged as potential mediators of RONS entry into plant cells, linking plasma exposure to downstream physiological effects.

AQPs are integral membrane proteins that facilitate the selective and passive transport of water and small solutes across cellular membranes in all forms of life, including plants [9–11]. In higher plants, the plasma membrane intrinsic protein (PIP) subfamily, particularly isoforms like PIP2;1, plays a crucial role in regulating water homeostasis, nutrient uptake and adaptation to environmental stresses [12, 13]. Each AQP monomer forms a distinct water-conducting channel composed of six transmembrane helices and conserved structural elements such as the Asn-Pro-Ala (NPA) motifs and the aromatic/arginine (ar/R) selectivity filter [14, 15]. In plants, additional regulation of AQP activity occurs via phosphorylation, pH gating and oxidation-sensitive features such as disulfide bonds between extracellular cysteine residues [16, 17]. While AQPs are well established as water channels, several isoforms have also been shown to facilitate the transport of small neutral solutes and, in some cases, hydrogen peroxide [18, 19]. Nonetheless, comprehensive computational studies on the ability of plant AQPs to conduct plasma-generated RONS are still lacking.

Earlier computational work by R Cordeiro [20] used molecular dynamics (MD) simulations to investigate the permeability of reactive oxygen species such as H_2_O, H_2_O_2_, HO_2_ and·OH through both mammalian and plant AQPs, revealing that small radicals may transiently access the pore. Yusupov et al. [21] extended this line of inquiry by modeling RONS interactions with AQP1 channels and lipid bilayers containing *plasma-induced oxidative modifications*, highlighting that membrane composition and oxidation state strongly influence permeation. However, these studies primarily focused mainly on mammalian AQP1 which is one of the important transmembrane protein in biomedical contexts and did not specifically address plant growth-related RONS transport.

In this work, we perform atomistic MD simulations [22, 23] to explore the permeability of plant aquaporin PIP2;1 to selected RONS (e.g., H_2_O_2_, HO_2_, NO, O_3_, OH, NO_2_, N_2_O_4_ HNO_3_, s-*cis*-HONO, and s-*trans*-HONO that are commonly generated by CAP. Our objective is to quantify the free energy barriers and structural interactions that govern RONS entry through AQPs, with the broader goal of identifying species most likely to penetrate plant cells and mediate growth-promoting effects. This mechanistic study represents a fundamental step within a larger agricultural project aimed at optimizing CAP-based technologies for enhanced crop performance. Understanding the molecular basis of RONS permeation through AQPs will inform the design of plasma devices optimized for enhanced nutrient uptake and stress tolerance in crops.

## Methods

We conducted atomistic MD simulations to investigate the transmembrane transport behavior of selected RONS through the plant AQP in a cell membrane. Particularly, the investigated RONS encompassed both hydrophilic species (HNO_3_, HO_2_, *cis*-HNO_2_, *trans*-HNO_2_) and hydrophobic species (NO, NO_2_, N_2_O_4_, and O_3_), covering a diverse range of chemical reactivity, polarity, and molecular size. These RONS were selected based on their relevance in environmental and plasma-treated agricultural systems, where they are known to interact with biological membranes. In addition, the molecular system comprised a tetrameric plant aquaporin (PIP2;1) obtained from the Protein Data Bank (PDB ID: 1Z98), embedded in a 1,2-dioleoyl-sn-glycero-3-phosphocholine (DOPC) lipid bilayer (see Fig. 1). The bilayer was solvated with explicit water layers on both the extracellular and cytoplasmic sides, mimicking the native plant cellular environment. Moreover, chloride ions (Cl^―^) were added to neutralize the system’s net charge. Additionally, the structural renderings of the systems were prepared using Visual Molecular Dynamics [24].

**Figure 1.**
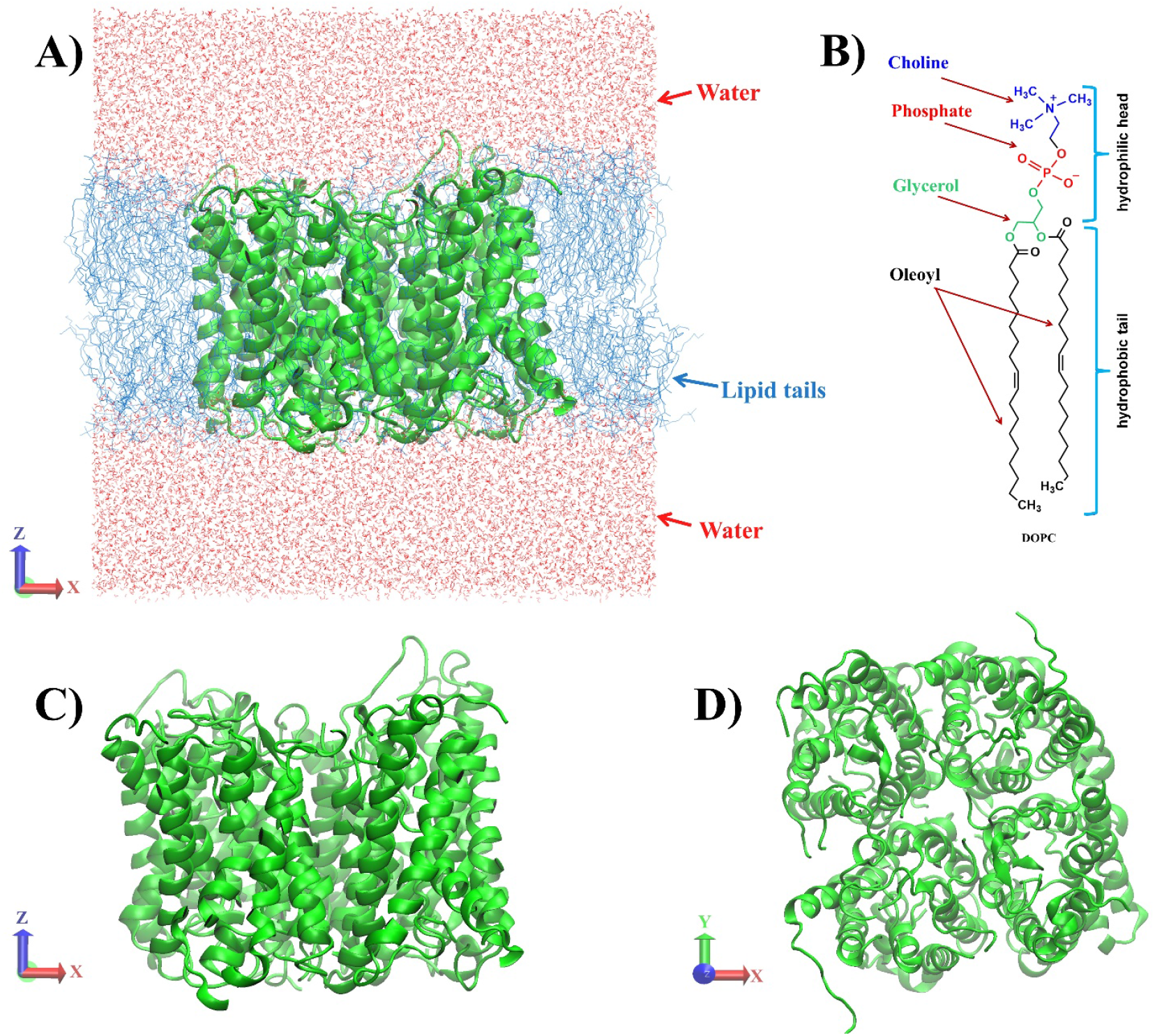
Structural configuration of the PIP2;1 aquaporin system: (A) side view showing the protein embedded in a DOPC lipid bilayer, (B) chemical structure of the DOPC lipid molecule, showing its amphipathic character, (C) side view of the tetrameric AQP arrangement, and (D) top view of the AQP

All MD simulations were carried out using GROMACS 2022 [25, 26]. The system was energy-minimized using the steepest descent algorithm for 50,000 steps, with a convergence criterion of 100 kJ·mol^−1^·nm^−1^. Long-range electrostatics were treated using the Particle Mesh Ewald (PME) method, and a 1.0 nm cutoff was applied for both Coulomb and van der Waals interactions. This was followed by NPT equilibration for 100 ps using a 0.5 fs time step. The system temperature was maintained at 298.15 K via the Nose-Hoover thermostat, and pressure was controlled semi-isotropically at 1.01325 bar using the Parrinello-Rahman barostat. Separate coupling groups were defined for the protein-lipid complex and the water-ion phase. Following equilibration, a 1 μs production MD simulation was conducted using a 2 fs timestep (total of 500 million steps). All thermostat and barostat parameters remained unchanged. The structural stability of the system was confirmed using root mean square deviation (RMSD) analysis of the protein backbone (see Fig. 3). A well-equilibrated frame from the final frame of this trajectory was selected as the starting configuration for umbrella sampling (US).

To probe the permeability of plasma-derived RONS through the AQP pore, US simulations were performed. At this stage, selected RONS, including N_2_O_4_, HNO_3_, HO_2_, *cis*-HNO_2_, *trans*-HNO_2_, NO, NO_2_, and O_3_, were individually inserted into the structure, which was extracted from the equilibrated system (see Fig. 2). RONS molecules were placed along the four individual pores of the AQP tetramer at 1.0 nm intervals along the z-axis (see Fig 2B). The inter-channel distance between solutes exceeded 1.0 nm, ensuring that each molecule remained beyond the cutoff radius for non-bonded interactions. As a result, no direct interactions occurred between RONS located in different channels. Each RONS was restrained in the z-direction using a harmonic potential (force constant = 2000 kJ·mol^−1^·nm^−2^) centered at the umbrella window. To limit lateral diffusion, a flat-bottomed restraint was applied in the xy-plane with a radius of 0.5 nm and a force constant of 500 kJ·mol^−1^·nm^−2^. The inter-solute distance exceeded the cutoff for non-bonded interactions, ensuring statistical independence of each solute’s sampling trajectory.

**Figure 2.**
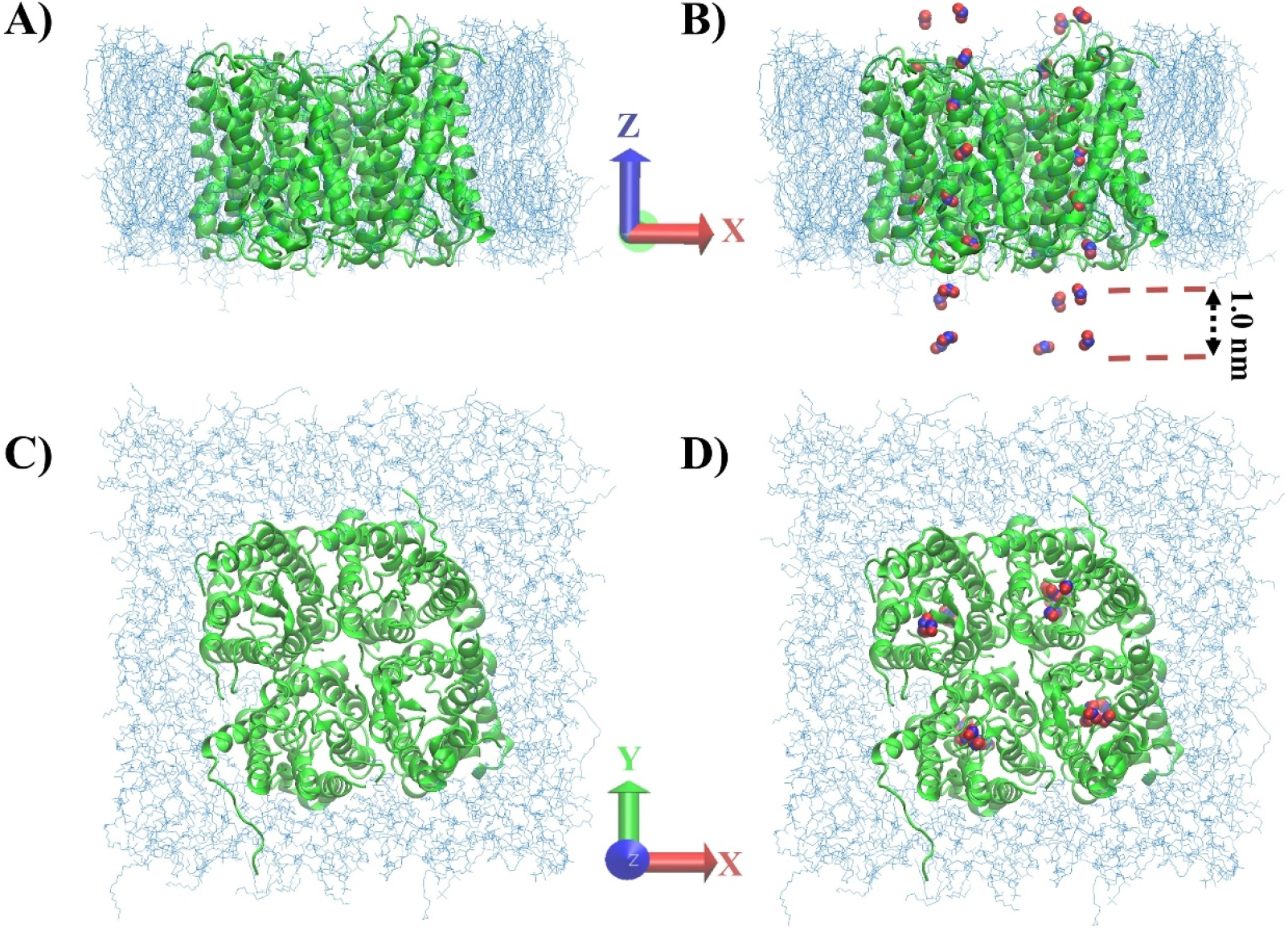
Preparation of umbrella sampling configurations: (A) side view of the equilibrated AQP-lipid system (water omitted for clarity), (B) side view after insertion of RONS along the z-axis, (C) top-down view of the tetrameric AQP system, and (D) top view showing RONS distribution within individual channels.

**Figure 3.**
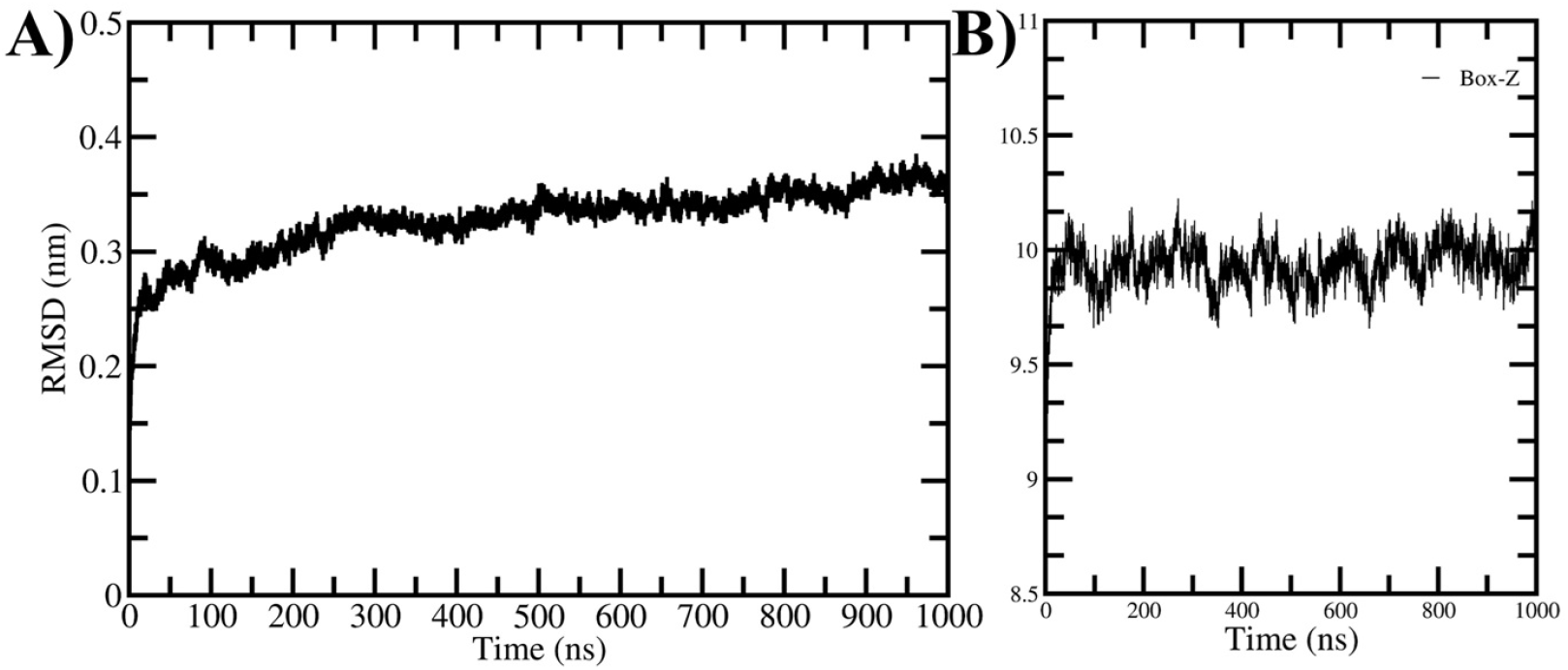
Global structural stability. (A) RMSD of PIP2;1 during the 1 µs MD simulation. (B) Box dimension along the Z-axis demonstrating stable bilayer thickness.

This parallelized US approach allowed simultaneous simulation of 32 RONS molecules per RONS type (8 per channel × 4 channels), optimizing computational efficiency while ensuring thorough sampling of free energy landscapes.

## Results and Discussion

To ensure reliable sampling of RONS permeation through the PIP2;1 aquaporin, a 1 μs atomistic MD simulation was conducted following standard minimization and equilibration protocols (see Methods). Structural stability of the channel was monitored via the RMSD of the protein backbone atoms relative to the initial structure (see Fig. 3A). The RMSD increased during the initial relaxation phase and reached a stable plateau of approximately ~0.32-0.34 nm after ~80 ns, remaining steady for the remainder of the trajectory. This stable behavior confirms that the aquaporin maintained its native fold and achieved dynamic equilibrium prior to umbrella sampling calculations.

In addition to RMSD, the stability of the membrane environment was monitored by tracking the box length along the Z-axis, which reflects the bilayer thickness (see Fig. 3B). The Z-dimension remained highly stable around ~10 nm over the entire 1-μs trajectory, with only small thermal fluctuations. This consistency confirms that the lipid bilayer preserved its structural integrity and that no artificial compression or expansion occurred during the production simulation.

Local flexibility of the protein was assessed using residue-wise RMSF analysis of the four PIP2;1 monomers (Fig. 4). The transmembrane helices remained highly stable, with fluctuations generally below 0.20 nm, consistent with a rigid pore architecture. In contrast, increased mobility was observed in the N-terminal region, in extracellular loops (around residues 60-75 and 140-160), and in the C-terminal tail (residues 270-280), where RMSF values reached up to 0.5-0.6 nm. These flexible regions correspond to non-helical segments typical of plant aquaporins and do not affect the structural stability of the conduction pathway.

**Figure 4.**
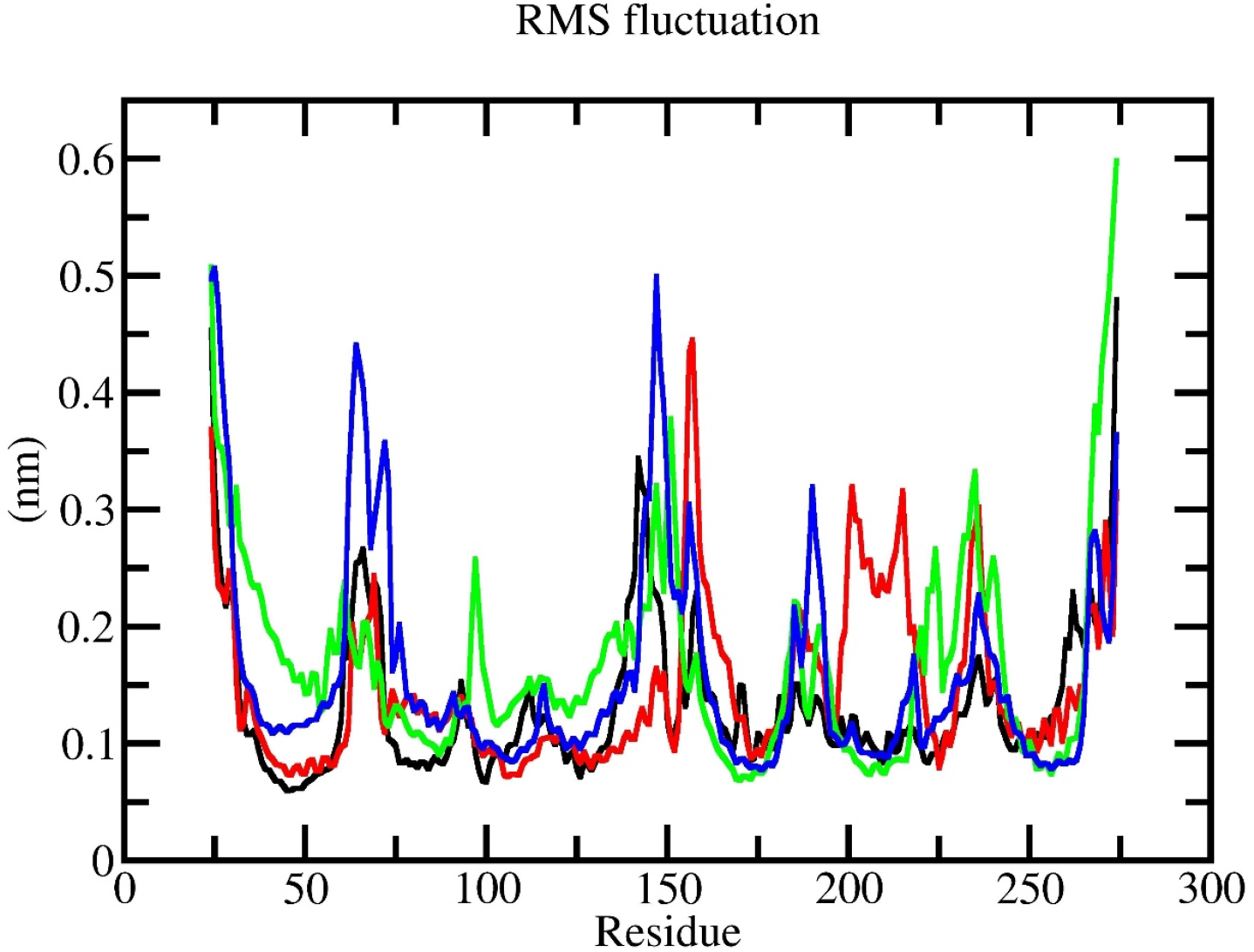
Residue-wise RMSF of PIP2;1, highlighting stable helices and flexible loop/terminal regions.

Figure 5 shows the free energy profiles (FEPs) for the relatively hydrophilic species - HO_2_, *cis*-HNO_2_, *trans*-HNO_2_, and HNO_3_ - calculated along the z-axis of the PIP2;1 channel. All four species were observed to enter the extracellular entry region of the channel with minimal energetic barriers but exhibited significantly different free energy barrier profile near the channel’s narrowest region, corresponding to the NPA motifs and ar/R filter. Among these, *trans*-HNO_2_ demonstrated the most favorable permeation profile, maintaining nearly barrierless translocation across the entire channel. The maximum free energy barrier along its path did not exceed ~2 kJ·mol^−1^, suggesting unhindered diffusion through the pore. In contrast, HNO_3_ displayed the highest free energy barrier for permeation, reaching approximately 13.6 kJ·mol^−1^ at the selective region. This is likely due to its stronger polarity and hydrogen bonding capacity, which result in stronger interactions with pore-lining residues and water molecules.

**Figure 5.**
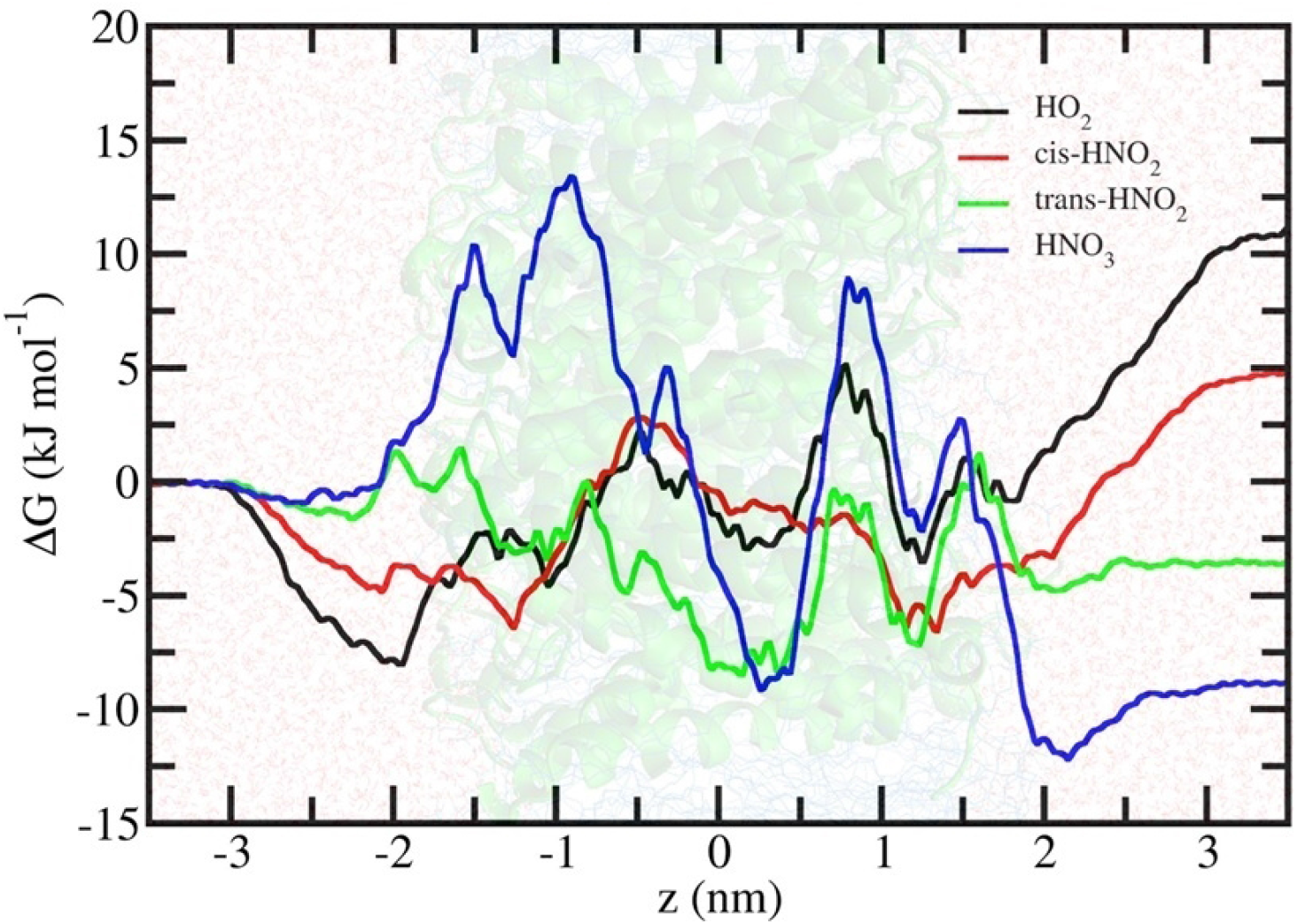
FEP of hydrophilic RONS - HO_2_, cis-HNO_2_, trans-HNO_2_, and HNO_3_ - along the z-axis of the PIP2;1 channel obtained from US simulations.

The FEPs of HO_2_ and *cis*-HNO_2_ presented intermediate behavior, with local barriers between 2-3 kJ·mol^−1^ near the pore center. The presence of multiple shallow minima along the permeation axis for HO_2_ indicates transient interactions with polar residues lining the pore. These results align with previous studies that suggested hydrophilic radicals may be sterically or electrostatically hindered in narrow AQP pores [20, 21]. The permeation FEPs for more hydrophobic or weakly polar species, NO, NO_2_, N_2_O_4_, and O_3_, are presented in Figure 6. Notably, NO exhibited a remarkably flat FEP with only minor barriers (~1-2 kJ·mol^−1^), suggesting near-free diffusion across the pore. This result is consistent with its small size, neutral charge, and minimal interaction with water or channel residues. O_3_ also displayed relatively favorable permeation, with barriers below 6 kJ·mol^−1^ and a smooth energetic landscape, supporting the hypothesis that weakly polar species can readily traverse narrow AQP pores. In contrast, N_2_O_4_ showed significantly higher barriers exceeding 15 kJ·mol^−1^ in the selective site, likely due to its bulkier geometry and dipolar character. NO_2_, which lies intermediate in polarity between NO and N_2_O_4_, exhibited moderate barriers (~6-12 kJ·mol^−1^), concentrated primarily at the channel center.

**Figure 6.**
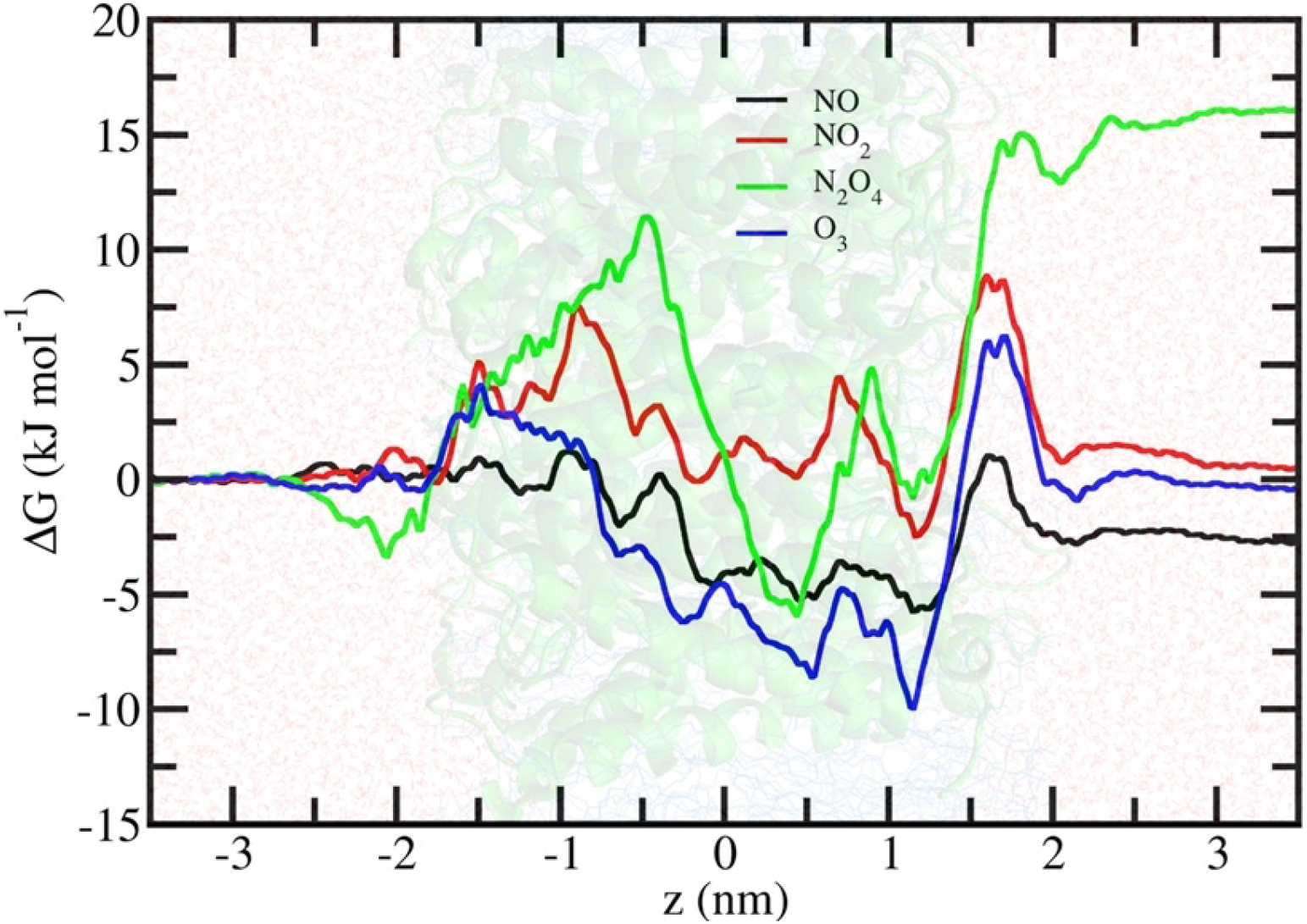
FEP of hydrophobic or weakly polar RONS - NO, NO_2_, N_2_O_4_, and O_3_ - across the PIP2;1 channel.

These findings highlight the dependence of RONS permeation on molecular size, polarity, and flexibility. Species with smaller van der Waals radii and weaker hydrogen bonding potential permeate more efficiently. Importantly, the energetics reported here correspond to passive, non-interacting transport; reactive events such as proton transfer, oxidation, or radical reactions are not captured within this framework but may further influence the biological impact of these species.

As summarized in Figure 7, the comparative analysis of free-energy barriers reveals a clear correlation between RONS polarity and permeability. The sequence of decreasing barriers (N_2_O_4_> HNO_3_> NO_2_ > O_3_ > HO_2_ ≈ cis-HNO_2_ > trans-HNO_2_ > NO) demonstrates that small, weakly polar species such as NO and trans-HNO_2_ diffuse most readily through PIP2;1, whereas bulky or highly polar molecules like HNO_3_ and N_2_O_4_ experience pronounced resistance at the ar/R constriction site.

**Figure 7.**
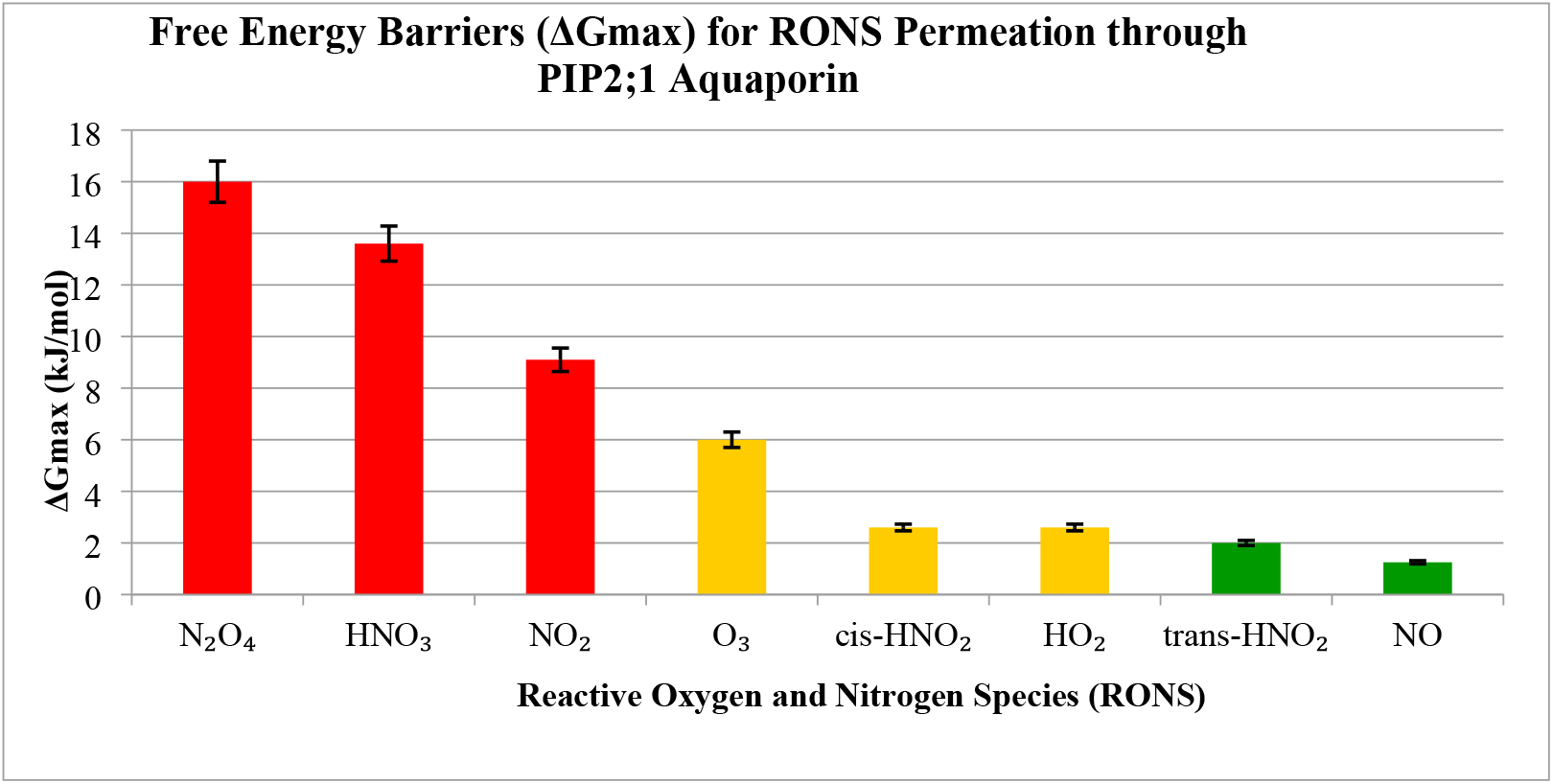
Comparative free-energy barriers (ΔG□_ax_) for plasma-derived reactive oxygen and nitrogen species (RONS) permeating through the plant aquaporin PIP2;1. Each bar represents the maximum free-energy barrier obtained from umbrella-sampling molecular-dynamics simulations. The color code highlights permeability levels: red - highly restricted species (HNO_3_, N_2_O_4_, NO_2_); gold - moderately permeable species (O_3_, HO_2_, cis-HNO_2_); and green - readily permeable species (trans-HNO_2_, NO). Lower ΔG_ax_ values correspond to easier diffusion through the aquaporin pore, indicating that NO and trans-HNO_2_ are the most favorable candidates for intracellular transport under plasma-treated agricultural conditions.

To further contextualize these results, the obtained ΔG□_ax_ values were compared with those previously reported for RONS permeation across lipid membranes without AQP [27]. While the absolute barriers differ due to structural and dielectric distinctions between lipid and protein channels, a similar trend is observed for polarity-dependent transport efficiency (Table 1). Notably, species such as NO and trans-HNO_2_ remain the most permeable in both systems, although the overall energy barriers in the PIP2;1 aquaporin are approximately 2-3 times higher owing to the tighter ar/R constriction. This comparison confirms that the AQP pore imposes stronger steric and electrostatic selectivity than lipid membranes, thereby serving as a highly discriminating gate for plasma-derived reactive species.

**Table 1.** Comparison of calculated maximum free-energy barriers (ΔG□_ax_) for selected plasma-derived RONS across nitro-oxidized lipid membranes [27] and the plant aquaporin PIP2;1 (this study). Higher barriers observed for polar species in the AQP highlight the selective nature of the ar/R filter.

| Species | $\Delta G_{\square_{ax}}$ (kJ·mol<sup>-1</sup>),<br>Membrane (2023) [27] | $\Delta G_{\square_{ax}}$ (kJ·mol<sup>-1</sup>),<br>Plant AQP (this work) |
| --- | --- | --- |
| HO<sub>2</sub> | 6.4 | 2.6 |
| cis-HNO<sub>2</sub> | -0.1 | 2.6 |
| trans-HNO<sub>2</sub> | 5.6 | 2.0 |
| HNO<sub>3</sub> | 7.55 | 13.6 |
| N<sub>2</sub>O<sub>4</sub> | 0.1 | 16 |
| NO | 0.5 | 1.25 |
| NO<sub>2</sub> | 0.9 | 9.1 |
| O<sub>3</sub> | 1.6 | 6.0 |

The differential permeability profiles observed among RONS provide important mechanistic insight into how CAP-generated species might interact with plant tissues at the molecular level. Hydrophobic and weakly polar species like NO and O_3_ can efficiently cross the cell membrane through AQP such as PIP2;1, potentially reaching intracellular compartments and modulating physiological processes such as growth or stress resistance.

In contrast, the entry of highly polar or reactive species (e.g., HNO_3_) may be kinetically or sterically hindered, limiting their biological activity unless facilitated by alternative mechanisms (e.g., transient pore expansion or lipid-mediated diffusion). These results suggest that the composition of plasma effluents could be tuned to favor the generation of species with high membrane permeability, thereby maximizing agricultural benefits such as early ripening or stress adaptation.

## Conclusion

This study provides a molecular-level investigation into the passive permeation of plasma-derived RONS through the plant aquaporin PIP2;1, using extensive atomistic molecular dynamics and US simulations. The results reveal that small, weakly polar species such as nitric oxide (NO) and ozone (O_3_) experience minimal energy barriers, enabling efficient translocation through the AQP channel. In contrast, larger or highly polar molecules like nitric acid (HNO_3_) and dinitrogen tetroxide (N_2_O_4_) face significant barriers near the selectivity filter, suggesting restricted passive permeation under physiological conditions.

These findings emphasize the importance of physicochemical properties, such as size, polarity, and hydrogen bonding capacity, in determining RONS transport efficiency through plant AQPs. The permeation profiles obtained here serve as a predictive framework for identifying which plasma-generated species are most likely to penetrate plant cells and exert downstream effects.

Importantly, this work contributes to a broader agricultural initiative aimed at optimizing CAP technologies for sustainable crop enhancement. By linking molecular permeability to potential intracellular activity, these results help guide the rational design of plasma treatments for improved nutrient uptake, stress tolerance, and accelerated growth in plants.

## Acknowledgments

J.R gratefully acknowledges the financial support received from the Agency for Innovative Development of the Republic of Uzbekistan, Grant Number FZ-2020092817.

## Data availability

The datasets used and/or analyzed during the current study are available from the corresponding author on reasonable request. All data generated during this study are included in this published article.

